# DNA methylation signature in *NSD2* loss-of-function variants appeared similar to that in Wolf-Hirschhorn syndrome

**DOI:** 10.1101/2023.01.06.522834

**Authors:** Tomoko Kawai, Shiori Kinoshita, Yuka Takayama, Eriko Onishi, Hiromi Kamura, Kazuaki Kojima, Hiroki Kikuchi, Miho Terao, Tohru Sugawara, Ohsuke Migita, Masayo Kagami, Tsuyoshi Isojima, Yu Yamaguchi, Keiko Wakui, Hirofumi Ohashi, Kenji Shimizu, Seiji Mizuno, Nobuhiko Okamoto, Yoshimitsu Fukushima, Fumio Takada, Kenjiro Kosaki, Shuji Takada, Hidenori Akutsu, Kiyoe Ura, Kazuhiko Nakabayashi, Kenichiro Hata

## Abstract

**Purpose:** Wolf-Hirschhorn syndrome (WHS), a contiguous gene syndrome caused by the hemizygous deletion of the distal short arm of chromosome 4 where *NSD2* is, reportedly exhibits specific DNA methylation signatures in peripheral blood cells. However, responsible genomic loci for signatures are unreported. The objective of the study is to define the loci of WHS-related DNA methylation signatures and to explore the role of *NSD2* for the signatures.

**Methods:** We conducted genome-wide methylation analysis of individuals with WHS or *NSD2* variants using array. We studied genome-edited knock in mice or induced pluripotent stem cells to explore the function of *NSD2* variants which are observed in congenital anomaly cases.

**Results:** Three undiagnosed cases with *NSD2* variants showed WHS-related DNA methylation signatures. These variants were validated to be *NSD2* loss-of-function in induced pluripotent stem cells or genome-edited knock-in mice. p.Pro905Leu variant decreased Nsd2 protein levels, and changed Histone H3-Lysine 36 demethylation levels in similar way in the same genomic regions as *Nsd2* knock out mice regulated. *Nsd2* knock out mice exhibited common DNA methylation changes.

**Conclusion:** These results revealed that WHS-related DNA methylation signatures are dependent on *NSD2* dysfunction and are useful in diagnosing *NSD2* variants of unknown significance.

## Introduction

Wolf-Hirschhorn syndrome (WHS, OMIM 194190), also known as 4p deletion syndrome, is a contiguous gene syndrome caused by the partial deletion of the short arm of chromosome 4 (4p) in a single allele. The critical region for phenotypes located on chromosome 4p16.3 contains 3 genes namely *LETM1* (OMIM *604407), *NSD2* (also named *WHSC1*, OMIM *602952), and *NELFA* (also named *WHSC2*, OMIM *606026).^1,2^ Single variants in the *NSD2* gene can cause mild phenotypes of WHS.^3-6^ NSD2, a member of the NSD family, is a Histone-Lysine 36 (H3K36) N-Methyltransferase.^7^ *NSD1* (OMIM *606681) variants cause Sotos syndrome 1 (OMIM 117550) and a specific DNA methylation signature consequently.^8^ Many epigenome-related gene variants exhibit unique combinations of DNA methylation changes defined as “episignature”.^9-11^ *NSD2* variants also exhibit DNA methylation signatures.^12^ However, the specific loci of DNA methylation changes for *NSD2* variants remain unknown. In this study, we define WHS-related DNA methylation signatures and determine whether *NSD2* loss of function established through functional assays of variants of uncertain significance (VUSs) can be classified as WHS-related DNA methylation signature.

## Materials and Methods

### Patients

A total of 21 patients with del(4)(p16) (Table S1), 3 patients with undiagnosed neurodevelopmental disorders with congenital anomalies (Table S2), and 3 patients with *NSD2* likely benign variants (Table S3) were recruited. The deleted regions at chromosome 4 in 16 out of the 21 patients with del(4)(p16) have been previously described (Table S1).^13,14^ Some of the patients were originally recruited to Initiative on Rare and Undiagnosed Diseases for diagnosis using exome sequencing.^15^ *NSD2* variant nomenclature refers to the NM_133330.2 transcript. Pathogenicity classification is based on the American College of Medical Genetics and Genomics/Association for Molecular Pathology (ACMG/AMP) guidelines.^16^

### DNA methylation

Genomic DNA was obtained from the peripheral blood of the patients and controls. DNA methylation data were obtained using the Illumina Infinium Methylation EPIC BeadChip array as previously described (Illumina, San Diego, CA, USA).^17^ Additional control methylation data were obtained from publicly available datasets taken from the Gene Expression Omnibus (GEO: GSE154566, GSE179759, GSE166503). Overall, we used 142 assays of unrelated individuals from the general population without clinically obvious neurodevelopmental phenotypes. For training and testing set, 106 and 36 assays of healthy individuals were used, respectively (Table S4). These numbers are 6.6 and 9 times more than the number of patients in each set. To distinguish *NSD2* variant-specific DNA methylation changes from those in *NSD1* variants, we referred 213 control data profiled using the Illumina Infinium HumanMethylation450 BeadChip (450K) array. (GEO: GSE36064, GSE42861 (non-smokers aged ˂ 50 y), GSE74432).

### Identification of WHS-related methylation probes

DNA methylation in the WHS and control blood samples in the training set was assessed through linear regression modeling using M-values, by applying logit transformation on the beta value. Peripheral blood cell composition was analyzed as previously described.^18^ The cell composition were input into the linear regression model as separate covariates. Beta value variance between groups were compared using var.test. Clustering of DNA methylation data was performed using the heatmap.2 function from the gplots package. Distances were measured using the Euclidean formula. Hierarchical cluster analysis was performed using “ward.D”. Support vector machine (SVM) modeling was performed to classify cases by WHS-related DNA methylation signature in a training set, using the e1071 R package (v.1.7-9). SVM decision values ranging between 0 and 1 in the testing set were converted to probability scores using Platt’s scaling.^19^ For comparison between our identified WHS- and Choufani et al. identified Sotos-related DNA methylation using the 450K (GSE74432),^8^ we compared 213 and 142 control data assessed using the 450K and EPIC (Table S4) arrays in GEO database, respectively, and selected 299,928 probes with a mean DNA methylation value difference of < 0.1 between two devices. Differentially methylated probes in each *NSD2* and *NSD1* defects were identified by the multiple linear regression model with Bonferroni multiple-test correcting moderated p values adjusted for blood cell types < 0.05 and delta beta > 0.1. To distinguish *NSD2* defects and *NSD1* defects and normal controls by DNA methylation values, we identified probes following threshold; differed more than 0.15 in absolute beta between *NSD2* defects and controls with < Bonferroni corrected p: 5e^-10^ and differed between *NSD2* defects and *NSD1* defects with < Bonferroni corrected p: 5e^-10^.

### Functional assay of *NSD2* VUS

Induced pluripotent stem cells (iPSCs) were generated from Case 2’s and healthy control’s peripheral blood cells as previously described.^20^ Genome-edited mice carrying *Nsd2* NM_001081102.2 transcript c.2717C>T substitution (*Nsd2*^wt/P906L^) and *Nsd2* knock out mice (*Nsd2*^wt/-^) were generated using the CRISPER/Cas9 systems as previously described.^21^ The guide RNA sequences are listed in Table S5. In mouse genome, *Nsd2* c.2717C>T; p.Pro906Leu is an orthologous substitution of *NSD2* c.2714C>T; p.Pro905Leu, which Case 3 harbors. Transient expression of *NSD2* VUS in HeLa cells was performed using FuGENE 6 (Promega, WI, USA). pCMV-3xFLAG-*NSD2* and pCMV-3xFLAG-*NSD2* c.2714C>T plasmids were constructed by VectorBuilder (IL, USA); 24 h post transfection, cells were lysed with AllPrep DNA/RNA Mini Kit (Qiagen, Hilden, Germany) or RIPA for assays as described below.

### Mouse phenotypes

Body weights were assessed to compare growth of each genotype mice. For comparison of intra-uterine growth, body weight of wild type (n = 14), *Nsd2*^wt/P906L^ (n = 21), and *Nsd2* ^P906L/P906L^ (n = 7) at embryonic day (E)18.5 were compared in some littermates with relative ratio. Ones of wild type (n = 9), *Nsd2*^wt/-^ (n = 13), and *Nsd2* ^-/-^ (n = 7) were also compared similarly. For comparison of postnatal growth, body weight of wild type (n = 18: male, n = 15: female), *Nsd2*^wt/P906L^ (n = 17: male, n = 18: female), and *Nsd2*^wt/-^ (n = 9: male, n = 10: female) at age 8wk were compared in each sex with relative ratio. Sample size estimation were performed with power.t.test of Rpackage.

### DNA methylation in mouse tissue

Genomic DNA was extracted from mouse tissues using the AllPrep DNA/RNA Mini Kit, followed by bisulfite treatment using the Zymo EZ DNA Methylation-Gold™ kit (Zymo Research, Irvine, CA, USA). Genome-wide DNA methylation was analyzed using the Infinium Mouse Methylation BeadChip (Illumina). Methylation data were acquired using the iScan system and processed using GenomeStudio 2.0 (Illumina). The background was corrected using the method provided by Illumina. We removed probes with detection P values > 0.01 in at least one sample and filtered probes located on the X, Y, and mitochondrial chromosomes. This yielded 265,725 autosomal probes from 22 assays. DNA methylation differences were assessed through linear regression modeling using beta values.

### RNA-seq

Total RNA was isolated from the peripheral blood cells treated with RNAlater, using the RiboPure™-Blood Kit (Ambion). RNA amplification was performed using the SMART-Seq® v4 Ultra® Low Input RNA Kit for Sequencing (TaKaRa) and cDNA libraries were prepared using Nextera XT DNA Sample Preparation Kit (Illumina). Sequencing was performed using NovaSeq (Illumina). The data were aligned to GRCh38 and transcript count was performed using Dragen (Illumina) while referring to the GRCh38.35 gtf files. In mouse tissue samples, total RNA was isolated from thymocytes using the AllPrep DNA/RNA Mini Kit. RNA-seq libraries were prepared using the NEBNext UltraII Directional RNA Library Prep Kit for Illumina (New England BioLabs, Ipswich, MA, USA). Sequencing was performed using HiSeq X Ten (Illumina). The data were aligned to GRCm38 mm10 using HISAT2-2.1.0, and transcript count was performed using Cufflinks 2.2.1. Significant changes in transcript expression were calculated using Cuffdiff.

### Chromatin Immunoprecipitation Sequencing (ChIP-Seq)

ChIP assay was performed using ChIP Reagents (Nippon Gene, Tokyo, Japan). Thymocytes (1×10^6^) were fixed with 1% formaldehyde for 5 min. Cells were resuspended in SDS lysis buffer, and the lysate was sonicated using the S220 Focused-ultrasonicator (Covaris, MA, USA), to fragment the chromatin. The chromatin was purified through centrifugation and immunoprecipitated using Dynabeads M-280 Sheep anti-mouse IgG (Veritas Life Sciences, PA, USA) conjugated with anti-H3K36me2 antibodies (ab176921; Abcam, Cambridge, UK) in 1× RIPA (150 mM) buffer with protease inhibitors, for 2 h at 4 ℃. The chromatin-bound beads were washed sequentially with 150 mM and 500 mM 1× RIPA, and TE buffers. Next, the chromatin-bound beads were incubated in ChIP direct elution buffer with proteinase K (200 µg/ml), overnight at 65 ℃. DNA was purified using AMPure XP beads (Beckman Coulter, CA, USA) as per manufacturer’s instructions. ChIP-Seq libraries were prepared using NEBNext ChIP-Seq Library Prep Master Mix Set and Multiplex Oligos for Illumina. Sequencing was performed using HiSeq 2500 (Illumina). The data were aligned to the GRCm38 mm10 reference genome using BWA v.0.7.17. Multiple mapped reads and PCR duplicates were removed. Generated Bam files were processed using deepTools to generate a coverage track bigwig. The coverage was calculated as the number of reads per 100 bp bin using the reads per kilobase per million mapped reads (RPKM) normalization. Differences between the mice samples were filtered by 1 RPKM.

### Western blot analysis

Western blot analysis was performed as previously described.^22^ Anti-Histone H3 K36me2 (ab176921), anti-H3 (ab1791), and anti-NSD2 (ab75359) antibodies were procured from Abcam; ab75359 is a monoclonal antibody to a fusion protein, corresponding to amino acids 1-647 of Human GST-NSD2. Anti-DDDDK-tag antibody was purchased for detection of FLAG from MBL (Tokyo, Japan). Anti-ACTB was purchased from Merck (Darmstadt, Germany). Mice thymocytes or iPSCs were lysed with RIPA buffer. Gel electrophoresis and membrane transfer were performed as per manufacturer’s protocol (BioRad, Hercules, CA, USA).

## Results

### DNA methylation signature in WHS

We analyzed genome-wide DNA methylation values using peripheral blood cells of patients with WHS (n = 16, Table S1) and healthy individuals as controls (n = 106), as a training set. We selected 280 probes which showed more than |0.2| methylation beta value difference compared with the controls and significant with Bonferroni multiple-testing corrected *p* value < 2.4e-16 after statistically adjusted for blood cell type compositions. Out of 280 probes, 278 were hypo-methylated in WHS (Table S6). Beta value variances of 122 and 27 out of the 280 probes were significantly greater and less in WHS, respectively. The 280 probes distinguished methylation related to WHS from those in control samples through hierarchical clustering (Fig. 1a). Cluster analysis using the 280 probes clearly distinguished WHS cases (n = 4) in the testing set, in a cluster separate from the testing set controls (n = 36) (Fig. 1b). An SVM model using the 280 probes indicated that the 4 WHS cases in the testing set exhibited a DNA methylation pattern similar to the 16 WHS cases in the training set (Fig. S1a). These results validated incidence of WHS-related DNA methylation signatures of the 280 probes. Enrichment at the intergenic region was highest in the 280 probes (Fig. S1b). An individual with deleted 4p16.2p15.31 that did not include *NSD2* was considered a control (Fig. 1b), and exhibited low scores similar to other controls in the SVM model (Fig. S1a). This suggests that WHS-related DNA methylation signatures could be caused by *NSD2* loss-of-function variants.

**Figure 1.**
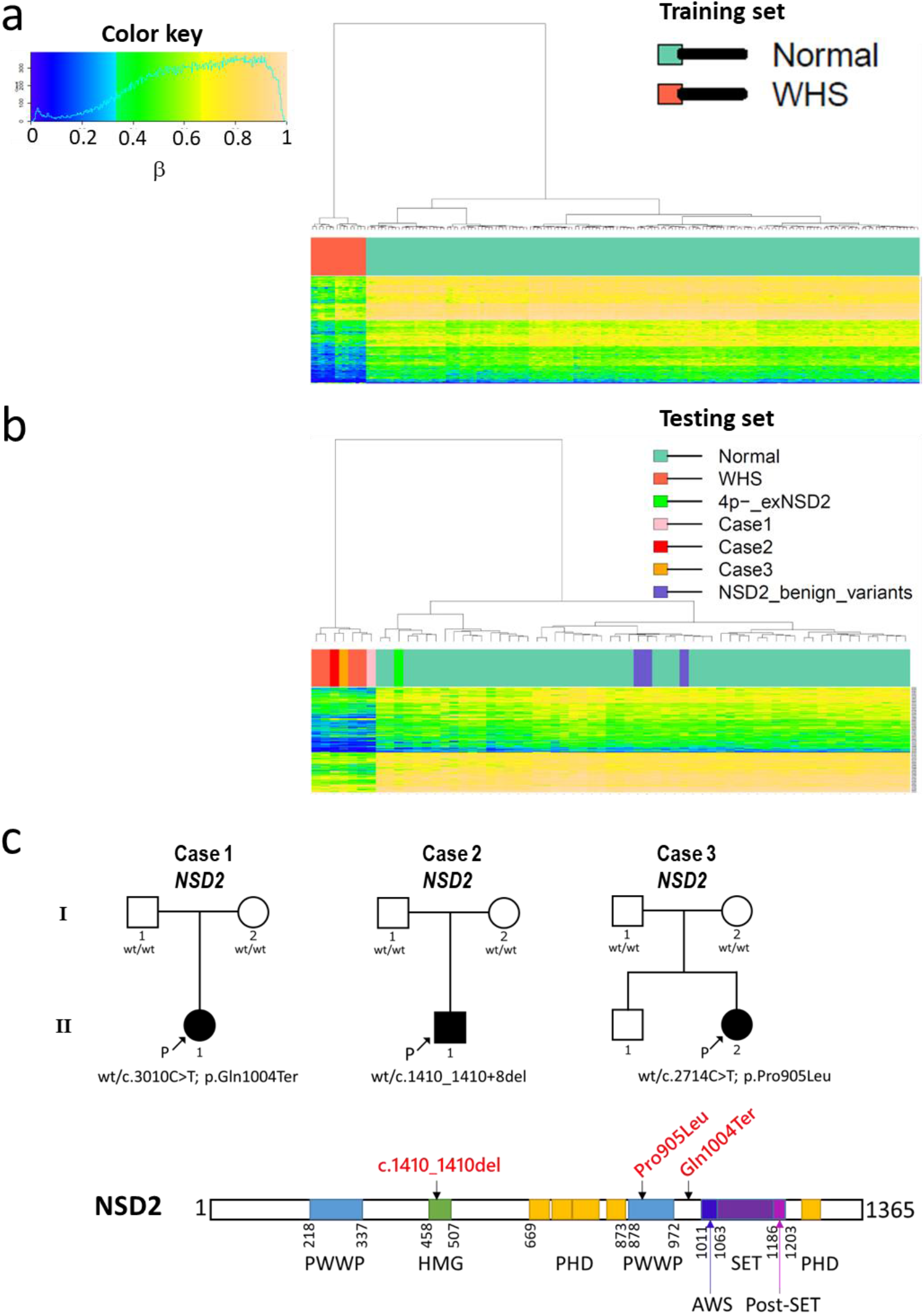
DNA methylation pattern of WHS and undiagnosed patients with *NSD2* variants. (**a**) The 280 probes identified in the training set by regression analysis were clearly separated into 16 WHS and 106 control samples through hierarchical clustering. (**b**) The 280 probes were validated to classify 4 WHS from 36 controls in the testing set. Three undiagnosed patients with *NSD2* VUSs were also classified into a WHS branch. Another three patients with likely benign *NSD2* variants were classified into a control branch. (**c**) Schematic illustration of variants of the 3 cases classified into a WHS branch in b.

### Evaluation of *NSD2* VUSs based on WHS-related DNA methylation signatures

We evaluated undiagnosed congenital anomaly cases with one *NSD2* “likely pathogenic” variant (Case 1), one *NSD2* “likely pathogenic” variant (Case 2), and one *NSD2* VUS (Case 3) (Table S2), based on the WHS-related DNA methylation signatures we established. Case 1 harbors a *de novo* nonsense variant with a single nucleotide substitution upstream of the *NSD2* SET domain (c.3010C>T; p.Gln1004Ter) (Fig. 1c). Case 2 harbors a *de novo* 9 bp deletion spanning exon 7 to intron 7 in *NSD2* (c.1410_1410+8del), which could cause abnormal splicing. Case 3 harbors a *de novo* missense variant in the highly conserved PWWP domain of *NSD2* (c.2714C>T; p.Pro905Leu). These three variants were clearly clustered in the same branch as WHS in the testing set and exhibited high scores in the SVM model (Fig. 1b and Fig. S1a). Meanwhile, the control cluster included three cases with “likely benign” *NSD2* variants (Fig. 1b and Table S3). These results strongly support that the WHS-related DNA methylation signatures isolated in this study could detect DNA methylation changes in undiagnosed cases with pathogenic *NSD2* variants.

### Functional assay of identified *NSD2* VUSs in undiagnosed patients

To validate that *NSD2* variants with WHS-related DNA methylation signatures are pathogenic, we conducted functional assays. Both expression of normal NSD2 proteins of approximately 150 kDa (O96028-1) and 75 kDa (O96028-3) and histone H3K36me2 levels, which is modified by NSD2, had decreased in Case 2 iPSCs, compared to control iPSCs (Fig. 2a and 2b). This indicates that c.1410_1410+8del in Case 2 is an *NSD2* loss-of-function variant. RNA-seq analysis of Case 2 peripheral blood cells revealed that c.1410_1410+8del in Case 2 is detectable as a transcript. The c.1410_1410+8del produced an abnormal transcript connected c.1410+464 at intron 7 to exon 8. The abnormal transcript bore premature termination codons and coded p.Glu470AspfsTer12, and escaped nonsense mediated decay (Fig. S2).

**Figure 2.**
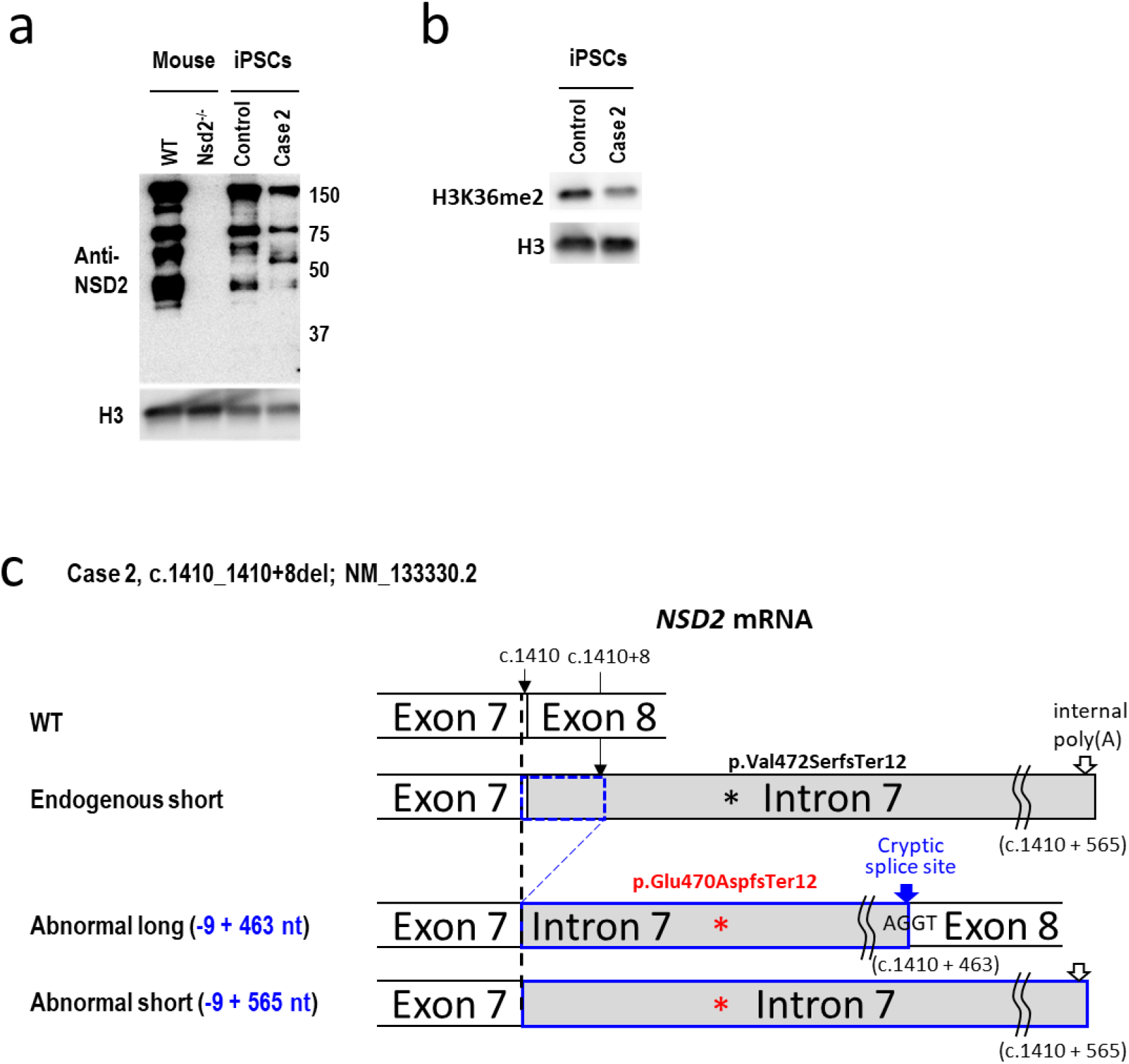
Effects of *NSD2* variants in Case 2. (**a**) *NSD2* antibody (ab75359) was specific to proteins derived from the Nsd2 protein via immunoblotting, since there was no band in *Nsd2*^*-/-*^ thymocytes. The decreased levels of NSD2 proteins (approx.150 kDa of O96028-1 and 75 KDa of O96028-3) were confirmed in the Case2 iPSCs. In contrast, a Case2-specific protein was detected by ab75359 (approx. 55kDa) which was close to the 53.18 kDa speculated molecular weight of NSD2 p.E470fs*12 by Expasy (https://www.expasy.org/). (**b**) The modification levels of H3K36me2 were decreased in Case2 iPSCs. (**c**) Schematic description of *NSD2* transcript from both alleles of Case 2.

Moreover, c.1410_1410+8del resulted in more usage of the internal poly(A) sequence (ENST00000508355.5) in intron 7 in *NSD2*. The resulting abnormal transcripts also code p.Glu470AspfsTer12 (Fig. 2c). Usage of the internal poly(A) sequence in *NSD2* was not detected in control samples (Fig. S2). Western blotting of Case 2 iPSCs with an NSD2 antibody detected an approximately 55 kDa Case 2-specific translated protein from these abnormal transcripts (Fig. 2a).

Since we did not reach a consensus on establishing Case 3 iPSCs, we generated knock-in mice carrying the mouse ortholog of Case 3’s *NSD2* variant i.e., *Nsd2*-p.Pro906Leu, and developed mice without the *Nsd2* coding region (Fig. 3a) that do not produce Nsd2 isoforms. The *Nsd2*-p.Pro906Leu knock-in homozygous mice (*Nsd2*^P906L/P906L^) rarely survived and all homozygous *Nsd2*-deleted mice (*Nsd2*^-/-^) exhibited embryonic lethality or died within a day after birth. Bodyweights of *Nsd2*^P906L/P906L^ and Nsd2^-/-^ mice were significantly lower than those of wild type (wt) and heterozygotic mice, respectively, at embryonic day (E)18.5. *Nsd2*^wt/P906L^ and *Nsd2*^wt/-^ mice were alive and fertile but exhibited significantly lower body weight than while type (wt) at age 8 weeks (Fig. 3b), which was concordant with Nimura et al.’s *Nsd2* mutant mice.^23^ Since *Nsd2* mRNA is highly expressed in the thymus^24^ and T cells are reportedly defected in *Nsd2* mutant mice, based on thymocyte analysis,^25^ we analyzed the fetal thymocytes^26^ of *Nsd2*^wt/P906L^ and *Nsd2*^P906L/P906L^ mice. Western blotting revealed decreased H3K36me2 in the *Nsd2*^wt/P906L^ and *Nsd2*^P906L/P906L^ mice, compared with wt mice (Fig. 3c). This indicates that the *NSD2* p.Pro905Leu is a loss-of-function variant. Moreover, *Nsd2*^P906L/P906L^ reduced Nsd2 expression (Fig. 3c).

**Figure 3.**
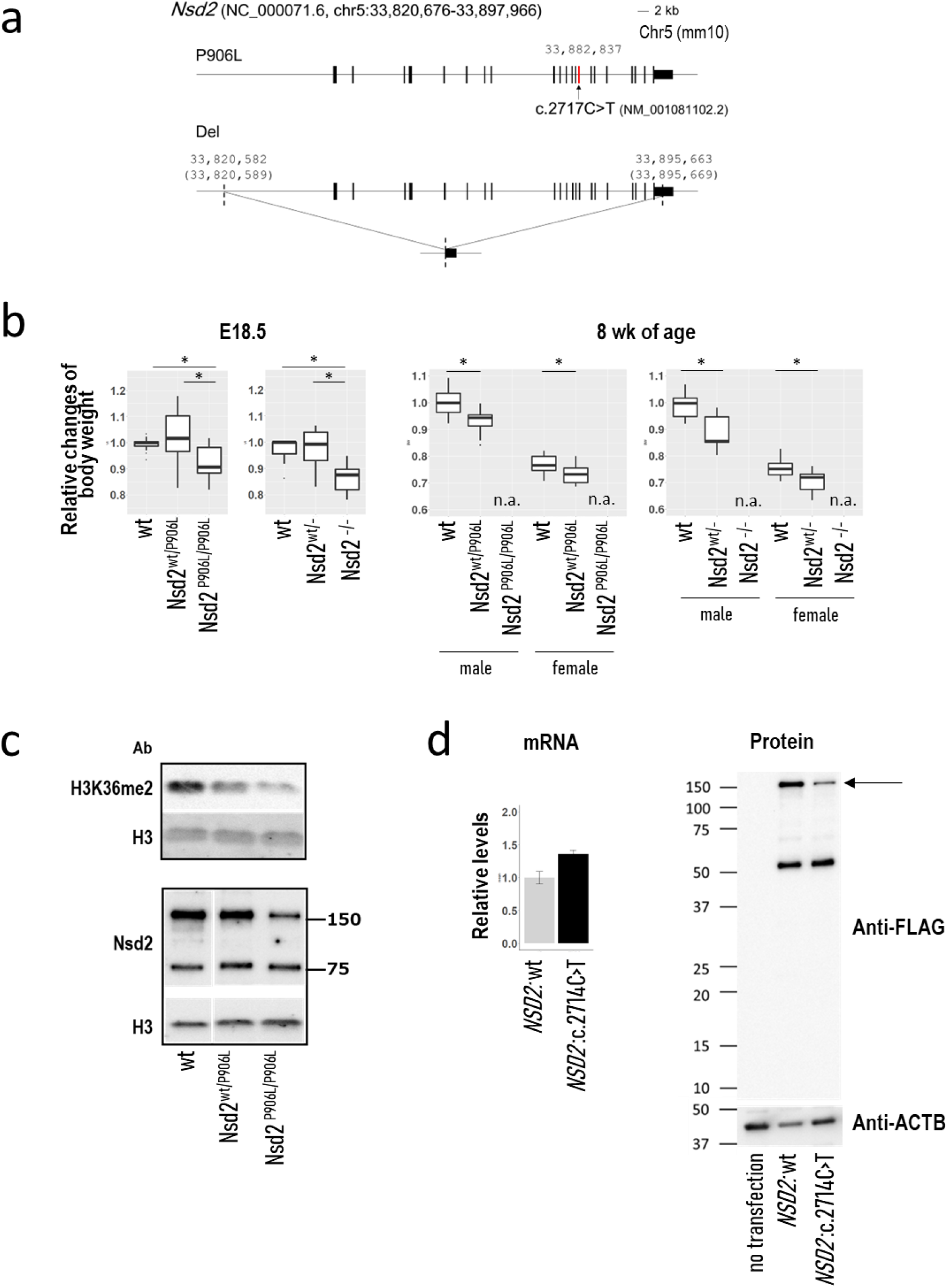
Functional assay of the Pro905Leu variant of NSD2. (**a**) Schematic description of *Nsd2*-variant mice used in this study. (**b**) Relative changes in bodyweight in *Nsd2-*variant mice, compared with the wild type mice at embryonic day (E)18.5 and 8 weeks of age. We showed relative changes since the number of littermates differed in mice by the embryonic lethality of the mutant mice. A significant decrease in intra-uterine growth was confirmed in homozygous mice of both variants. Failure to thrive was confirmed in heterozygous mice of both variants at 8 weeks of age. (**c**) *Nsd2*-p.Pro906Leu knock-in mice showed a decrease in H3K36me2 and Nsd2 protein in thymocytes at E18.5. (**d**) Overexpression of N-terminal FLAG-tagged *NSD2*: c.2714C>T in *NSD2* (NM_133330.2) coding p.Pro905Leu revealed a decrease in protein levels compared with the overexpression of N-terminal FLAG-tagged *NSD2* wild type. The arrow indicates approx.150 kDa of O96028-1.

Additionally, transient transfection of FLAG-tagged *NSD2* c.2714C>T; p.Pro905Leu exhibited decreased protein levels compared with transfection of FLAG-tagged *NSD2*, although mRNA expression levels were not changed between these (Fig. 3d). The decrease of protein levels was not associated with detection of extra bands with lower molecular weight or smeared band by Western blotting.

These results revealed that the *NSD2* VUS in Cases 1–3 was of the *NSD2* loss-of-function variants, showing that *NSD2* loss-of-function variants result in WHS-related DNA methylation signatures, as confirmed in Fig. 1b, which suggests that WHS-related DNA methylation signatures can be used to assess the pathogenicity of *NSD2* VUSs.

### *Nsd2*-p.Pro906Leu heterozygous knock in mice recapitulate the epigenetic changes in *Nsd2*^wt/-^ mice

We examined the effects of *NSD2* pathogenic variants on genome-wide H3K36me2 and comprehensive gene expression. ChIP-Seq of H3K36me2 in wt thymocyte-rich fractions showed broad genome-wide H3K36me2 modifications, similar to that in retinal pigment epithelial cells.^27^ Additionally, 62% of the H3K36me2 regions that exhibited more than 1 RPKM in 100 base-resolution were enriched in the genic region except for 5’UTR (Fig. 4a). Meanwhile, the number of H3K36me2-decreased regions (|> 1| delta RPKM) was the largest in the intergenic regions both in *Nsd2*^wt/-^ and *Nsd2*^wt/P906L^ thymocytes (Fig. 4b and 4c). Furthermore, increased H3K36me2 (> 1 RPKM) regions were observed both in *Nsd2*^wt/-^ and *Nsd2*^wt/P906L^ (Fig. 4c).

**Figure 4.**
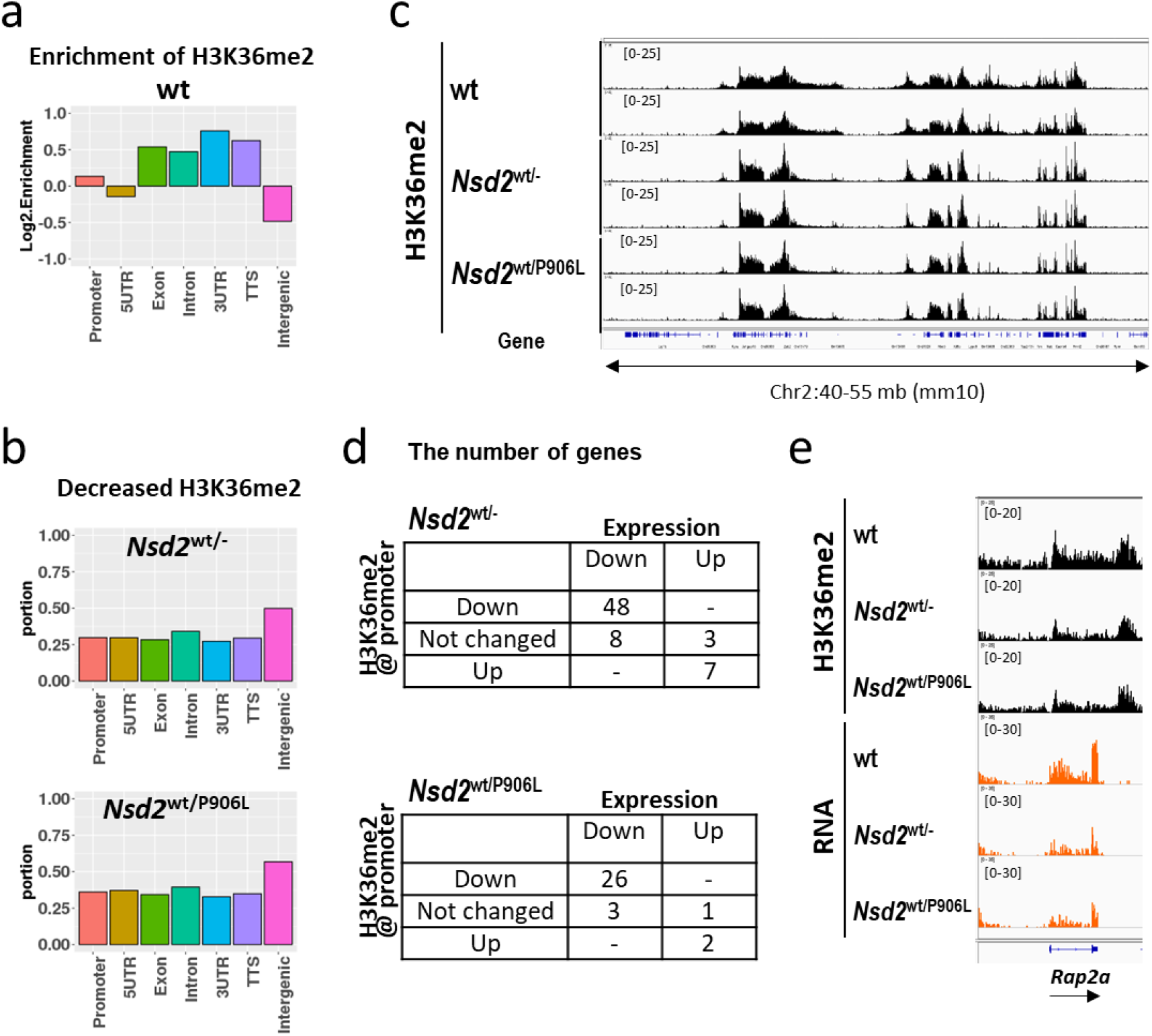
Multiomics analysis of *NSD2-*variants. (**a**) Genome-wide H3K36me2 enrichment by genetic features in wild type mice. (**b**) Proportion of H3K36me2-decrease windows by *Nsd2* KO or knock-in mice from wild type mice. (**c**) Representative genomic tracks of ChIP-Seq of H3K36me2 in each mice genotype. (**d**) Summary count of genes whose expression was regulated by variants. (**e**) Representative genomic tracks of ChIP-Seq of H3K36me2 and RNA-seq in each mice genotype.

H3K36me2 modification at the transcriptional start site (TSS) is correlated with gene expression levels when classified as High, Medium, and Low.^7^ The RNA-seq and ChIP-Seq of H3K36me2 confirmed that levels of H3K36me2 upstream of TSS were correlated to gene expression levels both in thymocytes and splenic B cells (Fig. S3a). *Nsd2*^wt/-^ thymocytes had 56 down-regulated and 10 up-regulated genes, compared with wt thymocytes (Fig. 4d). These expression changes were confirmed in genes expressed at “Medium” levels (Fig. S3b). Loss of H3K36me2 upstream of TSS was coordinated to 86% of down-regulated genes in *Nsd2*^wt/-^ thymocytes. Similar results were confirmed in *Nsd2*^wt/P906L^ thymocytes (Fig. 4d). Both *Nsd2*^wt/-^ and *Nsd2*^wt/P906L^ thymocytes down-regulated the same 21 genes, including *Rap2a* (Fig. 4e). These results suggest that *Nsd2* defects partially regulate the expression of genes, especially those expressed at “Medium” levels, at least in thymocytes. Additionaly, *NSD2*-p.Pro905Leu could coordinate gene expression in similar way with *NSD2* whole deletion through H3K36me2 changes.

### Concomitant changes between epigenetic changes and gene expression changes by *NSD2* defects

Next, to determine whether *NSD2* defects result in a DNA methylation signature, we performed a genome-wide analysis of DNA methylation of 265,725 CpGs in generated mice. We observed no significant differentially methylated CpG sites in *Nsd2*^wt/-^ and *Nsd2*^wt/P906L^ mice thymocytes, compared with wt mice thymocytes, at 8 weeks of age, which were identical thymocytes to that in ChIP-Seq and RNA-seq. DNA methylation changes were more noticeable in the few surviving *Nsd2*^P906L/P906L^ mice thymocytes compared with *Nsd2*^wt/-^ and *Nsd2*^wt/P906L^, at 8 weeks (Fig. S3c). Hence, we examined DNA methylation changes in accessible *Nsd2*^-/-^ fetal tissue. Significant hypo-methylation changes and common genome-wide DNA methylation changes in *Nsd2*^-/-^ were observed in fetal brain and heart (n = 5 each, Fig. S3d and S3e). These results validated that a single gene defect in *NSD2* can cause a DNA methylation signature.

To determine whether DNA methylation changes coordinate with gene expression changes, we assessed the DNA methylation changes of the CpGs at 1 kb ± TSS of the genes, whose expression were significantly down-regulated, in *Nsd2*^wt/-^ thymocytes, which were slightly hypermethylated (Fig. S3f). DNA methylation at the CpGs in the gene body of these genes were generally hypo-methylated in *Nsd2*^wt/-^ thymocytes. Hypo-methylation of CpGs in the gene body were also confirmed in the genes that are not expressed in thymocytes in a similar degree in *Nsd2*^wt/-^ thymocytes (Fig. S3g). It suggested that the gene expression changes from regional DNA methylation changes in the gene body included in *NSD2*-defects DNA methylation signature is not always the answer. Lastly, regarding to coordination between DNA methylation and H3K36me2 changes, concomitant changes with DNA hypo-methylation and H3K36me2 decrease were observed mostly in the intergenic region in *Nsd2*^wt/-^ mice (Fig. S3h). It may be one of the reasons that discrepancy between DNA methylation changes and gene expression changes. Overall, expression changes in *Nsd2*^wt/-^ thymocytes were more distinctly correlated to H3K36me2 changes than DNA methylation changes (Fig. S3i).

### DNA methylation differences in patients with *NSD2* loss-of-function variants and Sotos syndrome 1

NSD2 and NSD1 are H3K36me2 methyltransferases. The loss of function of *NSD2* and *NSD1* genes could cause similar epigenetic abnormalities. Indeed, patients with Sotos syndrome 1 exhibited high scores for WHS based on the SVM model using WHS-related DNA methylation signature (Fig. S1a and Table S7). However, as indicated by WHS and Sotos syndrome phenotypes, *NSD2* defects and *NSD1* defects influence development in different ways. Therefore, we attempted to identify DNA methylation changes in peripheral blood cells that would distinguish *NSD2* defects and *NSD1* defects and normal controls, using publicly available data. We identified 3,402 and 15,327 differentially methylated probes in *NSD2* and *NSD1* defects, respectively; 2,088 out of the 3,402 probes overlapped with the 15,327 probes. More robust changes in *NSD1* defects than in *NSD2* defects were confirmed (Fig. 5a). Only 20 probes clearly separated *NSD2* defects from *NSD1* defects and controls (Fig. S4a and Table S8). Among Sotos syndrome 1, Tatton-Brown Rahman syndrome with heterozygous *DMNT3A* (OMIM *602769) loss-of-function (TBRS, OMIM615879), immunodeficiency, centromeric region instability, facial anomalies syndrome 1 with homozygous *DMNT3B* (OMIM *602900) loss-of-function (ICF1, OMIM 242860), and *NSD2* defects, which are all characterized via genome-wide DNA hypomethylation,^10^ Sotos syndrome 1 showed unique DNA hypomethylation pattern (Fig. S4b). DNA methylation signature in *NSD2* defects were similar within these 4 syndromes (Fig. S4c). These results suggest that epigenetic regulation by *NSD2* were shared with other epigenetic regulators. To distinguish *NSD2* defects from other gene’s defects by DNA methylation signature, we confirmed whether the 856 probes, which Aref-Eshghi et al.^10^ reported to distinguish the 14 syndromes when the dimension of DNA methylation data were reduced, could cluster *NSD2* defects. The 856 probes clearly distinguished syndromes other than the 14 syndromes, but did not distinguish *NSD2* defects from controls (Fig. S4c). Adding the 280 probes of WHS-related DNA methylation signature to the 856 probes clearly distinguished *NSD2* defects from controls with no effects on classification of other syndromes (Fig. S4d). Finally, we confirmed that the top 50 probes of the 280 probes by p-value were sufficient to separate *NSD2* defects from controls and other syndromes by addition to the 856 probes (Fig. 5b).

**Figure 5.**
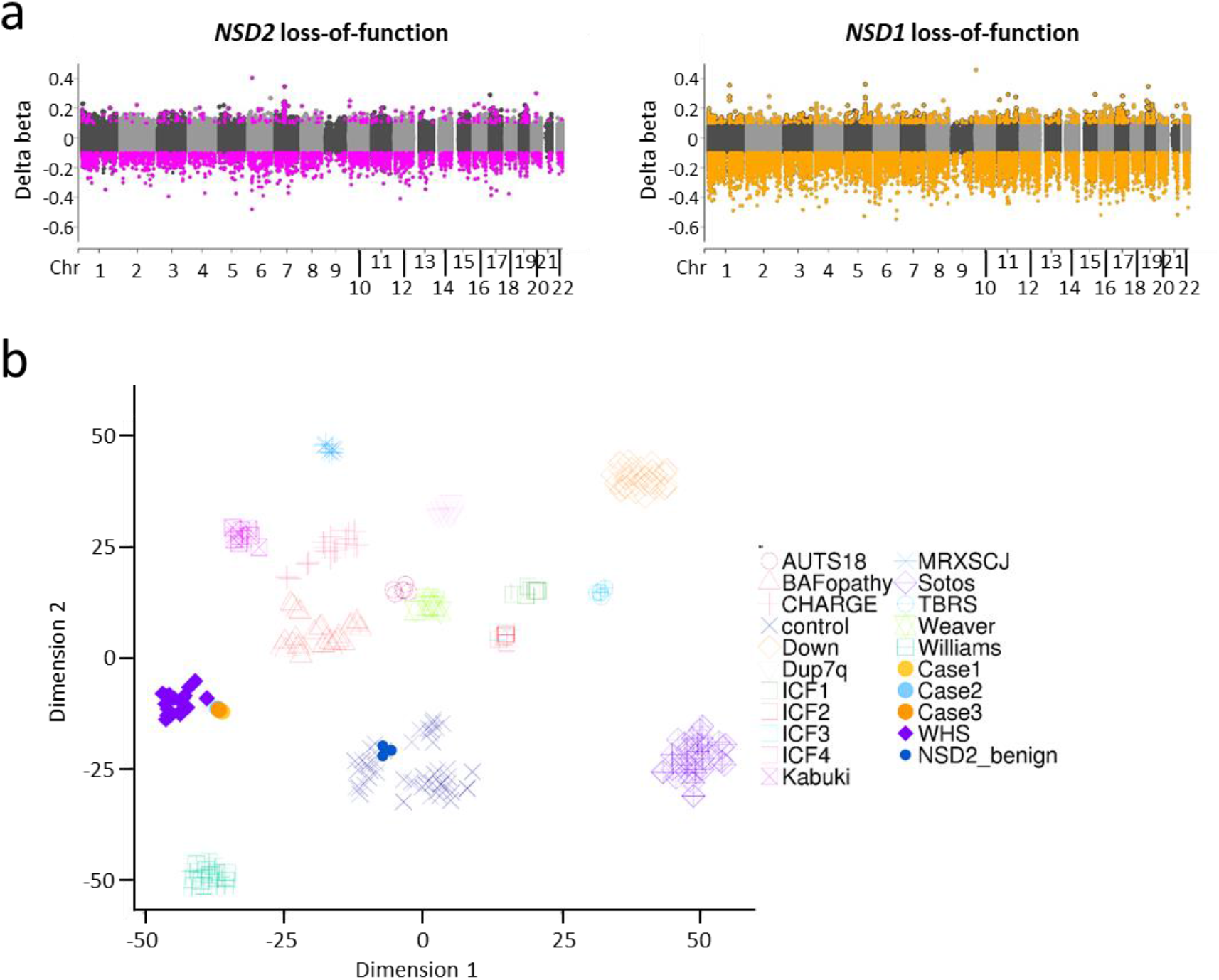
DNA methylation changes in *NSD2* loss-of-function variants and other syndromes. (**a**) Comparison of genome-wide DNA methylation changes between *NSD2* and *NSD1* loss-of-function variants in peripheral blood cells. Pink and orange dots indicate 3,402 and 15,327 probes with confirmed significantly differential methylation in *NSD2* and *NSD1* loss-of-function variants, respectively, with Bonferroni-corrected *p* < 0.05 and |delta beta| > 0.1. (**b**) Dimension reduction of DNA methylation data of subjects with WHS, *NSD2* loss-of-function variants, *NSD2* benign variants, and other syndromes for which “episignatures” have been reported. The cases are projected in two-dimensional plots using t-distributed stochastic neighbor embedding. Abbreviations: AUTS18: autism, susceptibility to, 18 (OMIM 615032); BAFopathy: Coffin-Siris 1,3,4 (OMIM 135900, 614608, 614609) and Nicolaides-Baraitser syndromes (OMIM 601358); CHARGE: CHARGE syndrome (OMIM 214800); Down: Down syndrome (OMIM 190685); Dup7q: Chr7q11.23 duplication syndrome (OMIM 609757); ICF1 to 4: immunodeficiency-centromeric instability facial anomalies syndrome1 to 4 (OMIM 242860, 614069, 616910, 616911); Kabuki: Kabuki syndrome 1 and 2 (OMIM 147920, 300867); MRXSCJ: Mental retardation, X-linked syndromic, Claes-Jensen type (OMIM 300534); TBRS: Tatton-Brown-Rahman syndrome (OMIM 615879); Weaver: Weaver syndrome (OMIM 277590); Williams: Williams syndrome (OMIM 194050).

## Discussion

Our results suggest that the WHS-related DNA methylation signatures identified in this study include the DNA methylation signature in *NSD2* loss-of-function variants and are useful in diagnosing the pathogenicity of *NSD2* VUSs. Although the DNA methylation patterns of the 3 cases with *NSD2* variants were similar to the WHS-related DNA methylation signature, these cases were not clinically diagnosed as WHS. Reportedly, *NSD2* loss-of-function variants lead to a distinct, rather mild phenotype partially overlapping with WHS.^3-6^ The overlapping phenotype could be related to common epigenetic dysfunctions like that in DNA methylation signatures.

H3K36me2 modifications were enriched in the genic region in wt mice; however, more decreased H3K36me2 modifications were observed in the intergenic region in adult *Nsd2* heterozygous KO and knock-in (i.e., *Nsd2*^wt/P906L^) mice thymocytes. This was consistent with Weinberg et al.’s findings for *Nsd1/2* genetically ablated mesenchymal stem cells and *Nsd1* genetically ablated embryonic stem cells.^28^ Increased H3K36me2 was also detected in some genomic regions in *Nsd2*^wt/-^ and *Nsd2*^wt/P906L^ mice. This conflicting phenomenon was also reported in multiple myelomas with *NSD2* variants^7^ or squamous cell carcinomas with *NSD1* variants.^29^ It might be a result of the redundancy of other enzymes for H3K36me2 modification than NSD2 and/or the balance between reciprocal modification of H3K36me2 and H3K27me3^30^ that also involves Histone H1.^31,32^ Most down-regulated genes in *Nsd2*^wt/-^ and *Nsd2*^wt/P906L^ mice correlated with decreased H3K36me2 before TSS (Fig. S3b). *NSD2* is reportedly located near TSS but avoids it,^33^ and involved in transcription.^34^ These genes could be involved in pathological conditions as direct targets of *Nsd2* loss-of-function variants. Although there were no differentially methylated loci in *Nsd2*^wt/-^ and *Nsd2*^wt/P906L^ mice in a statistically significant manner, hyper-methylation around TSS and general hypo-methylation across the entire gene body of significantly down-regulated genes were observed (Fig. S3f, g). However, DNA methylation changes occurred at similar levels in the gene body of non-expressed genes (Fig. S3g). These results highlight the difficulty in speculating gene expression changes through DNA methylation changes in the gene body. The effects of DNA methylation changes on gene expression could be related to tissue-specificity of gene expression.^35^

H3K36me2 recruits DNMT3A.^28,36,37^ DNA hypomethylation were most often observed in the intergenic region, along with H3K36me2 decrease caused by *Nsd2* variants (Fig. S3h) as previously reported,^28^ and in blood cells of patients with *NSD2* defects (Fig. S1b). However, DNA methylation is regulated by other histone modifications, too.^38^ DNA methylation changes were observed not only in regions that H3K36me2 was regulated by *Nsd2* variants but also in regions where H3K36me2 were undetected in all genotypes (Fig. S3c). It is possible that the DNA methylation signature identified herein, specific to *NSD2* loss-of-function variants, might not be direct targets of H3K36me2 by *NSD2*. In contrast, episignature is reported in each single epigenetic regulator gene’s variants.^11^ We distinguished patients with *NSD2* loss-of-function variants from other syndromes using WHS-related DNA methylation signatures (Fig. 5b). This indicates common epigenetic and etiological backgrounds in patients with *NSD2* loss-of-function variants.

Our study revealed the DNA methylation signature in *NSD2* loss-of-function variants and *NSD2*-p.Pro905Leu as a loss-of-function variant. Opposite DNA methylation patterns between Sotos syndrome 1 and Hunter McAlpine syndrome with duplication of *NSD1* (OMIM 601379) compared with controls have been previously reported.^10,39^ This indicates that opposite DNA methylation patterns could be identified between *NSD2* gain-of-function and loss-of-function variants. In addition, distinct DNA methylation signatures in Helsmoortel-van der Aa Syndrome (OMIM 615873) have been reported based on the loci of missense variants in *ADNP* gene (OMIM *611386).^40^ Regarding *KAT6B* (OMIM *605880), variants resulted in distinct DNA methylation signatures are associated with different syndromes—namely, Genitopatellar syndrome (OMIM 606170) and Say-Barber-Biesecker-Young-Simpson syndrome (OMIM 603736).^10^ Hence, DNA methylation patterns of the 280 probes could be criteria to assess of function of *NSD2* variants.

## Data Availability

All DNA methylation array and sequencing data have been deposited in the Gene Expression Omnibus (GEO) at **GSE174251** (human methylation array), **GSE176070** (RNA-seq), **GSE220890** (ChIP-Seq), and **GSE221621** (mouse methylation array).

## Acknowledgements

English language editing services were provided by Editage.

## Funding Statement

This work has been funded by research grants from the Japan Agency for Medical Research and Development (AMED) (Practical Research Project for Rare/Intractable Diseases; ek0109489 to K.H., ek0109205 to T.K.), JSPS KAKENHI (21H02887 and 21K19584 to K.H.), National Center for Child Health and Development (NCCHD 2022A-3 to K.H.), Gunma university for the promotion of scientific research to K.H..

## Author Contributions

Conceptualization: T.K., K.H.; Data curation: K.N.; Formal analysis: T.K.; Funding acquisition: T.K., K.H.; Investigation: T.K., S.K., Y.T., E.O., K.K., H.K., M.T., T.S.; Methodology: S.T., H.A., K.U., K.N.; Resources: O.M., M.K., T.I., Y.Y., K.W., H.O., K.S., S.M., N.O., Y.F., F.T., K.K.; Visualization: T.K., S.K.; Writing-original draft: T.K.; Writing-review & editing; K.N., K.H.

## Ethics Declaration

Approval for review and reporting these cases was given by all the relevant Japanese medical institutions: National Center for Child Health and Development, St. Marianna University School of Medicine, Gunma Children’s Medical Center, Keio University School of Medicine, Kitasato University School of Medicine, Shinshu University School of Medicine, Osaka Women’s and Children’s Hospital, Saitama Children’s Medical Center, and Central Hospital, Aichi Human Service Center. Written informed consent was obtained from the individuals’ parents for genetic testing. Human cells were collected post ethical approval from the Institutional Review Board of National Institute for Child Health and Development, Japan. Signed informed consent was obtained from donors or their parents, and the specimens were irreversibly de-identified. All experiments were performed according to the tenets of the Declaration of Helsinki. All protocols for animal experiments were approved by the Animal Care and Use Committee of the National Research Institute for Child Health and Development, Tokyo, Japan.

## Conflicts of Interest

The authors declare no conflict of interest.

## Notes

### Competing Interest Statement

The authors have declared no competing interest.

