## Supplementary Figure for "DNA methylation signature in *NSD2* loss-of-function variants appeared similar to that in Wolf-Hirschhorn syndrome"

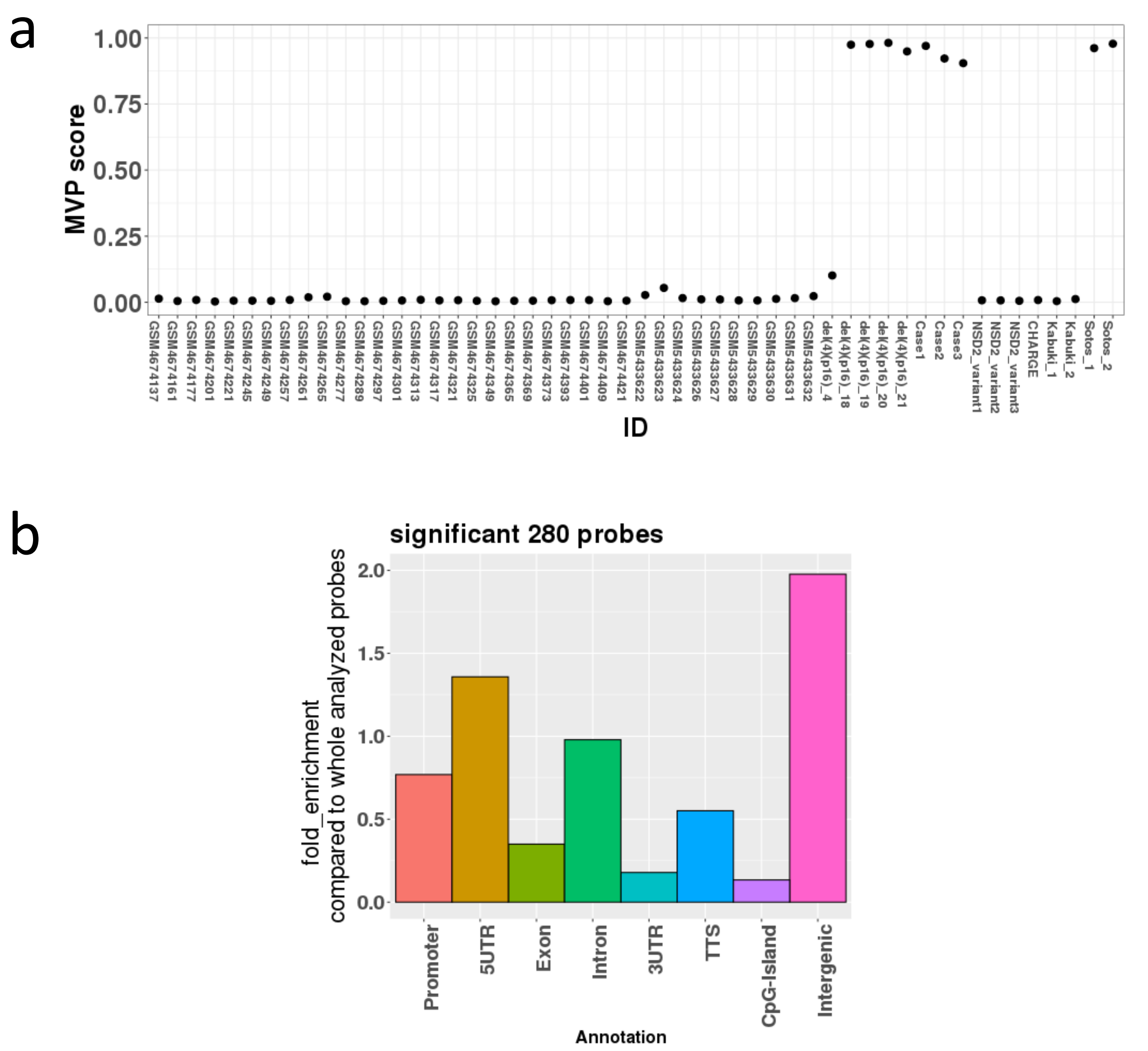


Figure S1. The identified 280 probes. (a) An SVM model using beta value of the 280 probes indicated that the 4 WHS cases (del(4)(p16)_18,19,20,21) in the testing set exhibited a DNA methylation pattern similar to the 16 WHS cases in the training set with high MVP scores. An individual with deleted 4p16.2p15.31 that did not include *NSD2* (del(4)(p16)_4) showed low MVP score. The 3 cases with *NSD2* variants (Case 1,2,3) showed high MVP score, while 3 cases with *NSD2* likely benign variants (NSD2_variant 1,2,3) showed low MVP score. (b) Genomic distribution of the 280 probes indicated an enrichment in 'Intergenic'.

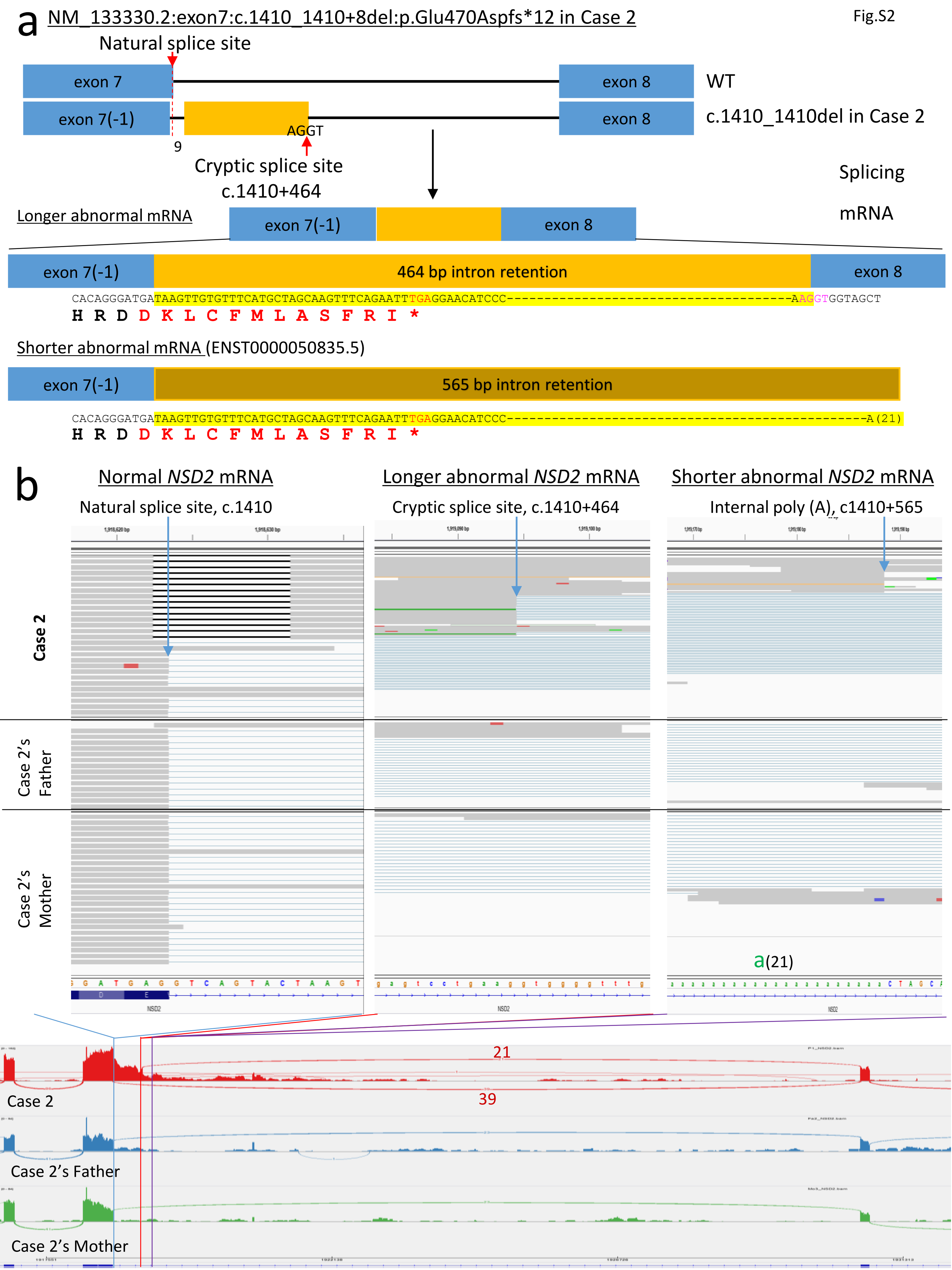
a
NM_133330.2:exon7:c.1410_1410+8del:p.Glu470Aspfs*12 in Case 2
Natural splice site
exon 7
exon 8
WT
exon 7(-1)
AGGT
exon 8
c.1410_1410del in Case 2
9
Cryptic splice site
c.1410+464
Splicing
mRNA
Longer abnormal mRNA
exon 7(-1)
exon 8
exon 7(-1)
464 bp intron retention
exon 8
CACAGGGATGATAAGTTGTGTTTCATGCTAGCAAGTTTCAGAATTTGAGGAACATCCC------------------------------------AAGGTGGTAGCT
H R D D K L C F M L A S F R I *
Shorter abnormal mRNA (ENST0000050835.5)
exon 7(-1)
565 bp intron retention
CACAGGGATGATAAGTTGTGTTTCATGCTAGCAAGTTTCAGAATTTGAGGAACATCCC--------------------------------------------------A(21)
H R D D K L C F M L A S F R I *
b
Normal NSD2 mRNA
Longer abnormal NSD2 mRNA
Shorter abnormal NSD2 mRNA
Natural splice site, c.1410
Cryptic splice site, c.1410+464
Internal poly (A), c1410+565
Case 2
Case 2's Father
Case 2's Mother
a(21)
21
39
Case 2
Case 2's Father
Case 2's Mother

Figure S2. Schematic description of *NSD2* transcript from *NSD2*_c.1410_1410+8del. (a) RNA-Seq of peripheral blood cells of Case 2 reveal a cryptic splice site in *NSD2* mRNA. Blue and yellow squares indicate exons and introns, respectively. (b) RNA-Seq of peripheral blood cells of Case 2 revealed existence of *NSD2* transcript with c.1410_1410+8del which indicated escape of nonsense mediated decay. Usage of the cryptic splice site was confirmed by some sequence reads. Case 2-specific transcript ends were confirmed at internal poly(A) site in intron 7 of *NSD2.*

Fig.S3

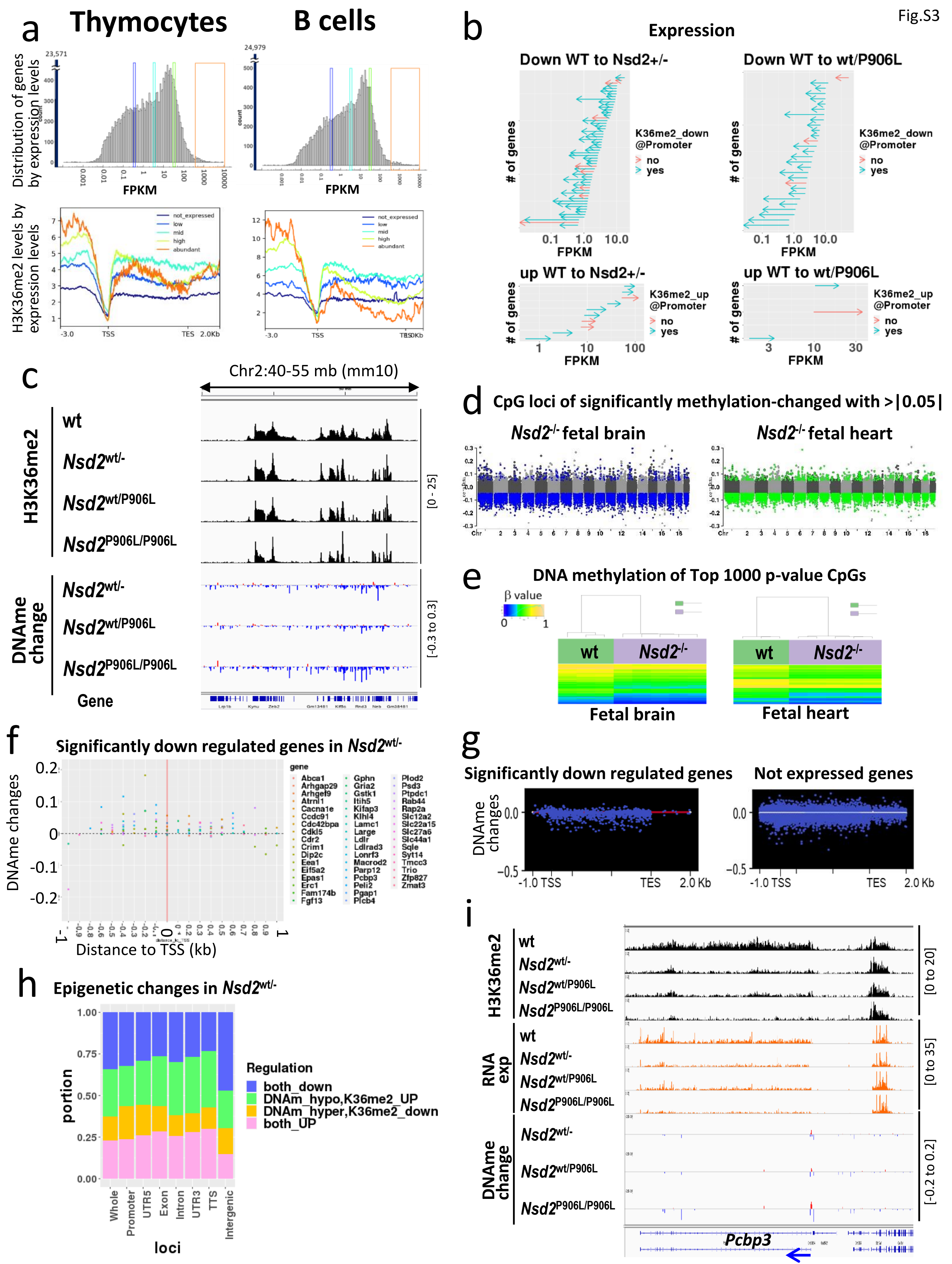

a
Thymocytes
B cells
Distribution of genes by expression levels
FPKM
H3K36me2 levels by expression levels
not_expressed
low
mid
high
abundant
TSS
TES
b
Expression
Down WT to Nsd2+/-
Down WT to wt/P906L
### of genes
FPKM
K36me2_down @Promoter
no
yes
up WT to Nsd2+/-
up WT to wt/P906L
K36me2_up @Promoter
c
Chr2:40-55 mb (mm10)
H3K36me2
wt
Nsd2wt/-
Nsd2wt/P906L
Nsd2P906L/P906L
[0 - 25]
DNAme change
[-0.3 to 0.3]
Gene
d
CpG loci of significantly methylation-changed with >|0.05|
Nsd2-/- fetal brain
Nsd2-/- fetal heart
e
DNA methylation of Top 1000 p-value CpGs
β value
wt
Nsd2-/-
Fetal brain
Fetal heart
f
Significantly down regulated genes in Nsd2wt/-
DNAme changes
Distance to TSS (kb)
gene
Abca1
Arhgap29
Arhgef9
Atrnl1
Cacna1e
Ccdc91
Cdc42bpa
Cdkl5
Cdr2
Crim1
Dip2c
Eea1
Eif5a2
Epas1
Erc1
Fam174b
Fgf13
Gphn
Gria2
Gstk1
Itih5
Kifap3
Klhl4
Lamc1
Large
Ldlr
Ldlrad3
Lonrf3
Macrod2
Parp12
Pcbp3
Peli2
Pgap1
Plcb4
Plod2
Psd3
Ptpdc1
Rab44
Rap2a
Slc12a2
Slc22a15
Slc27a6
Slc44a1
Sqle
Syt14
Tmcc3
Trio
Zfp827
Zmat3
g
Significantly down regulated genes
Not expressed genes
DNAme changes
-1.0 TSS
TES
2.0 Kb
h
Epigenetic changes in Nsd2wt/-
portion
Regulation
both_down
DNAm_hypo,K36me2_UP
DNAm_hyper,K36me2_down
both_UP
Whole
Promoter
UTR5
Exon
Intron
UTR3
TTS
Intergenic
loci
i
H3K36me2
RNA exp
DNAme change
[0 to 20]
[0 to 35]
[-0.2 to 0.2]
Pcbp3

Figure S3. Functional assay of *Nsd2* mutant mice. (a) Relevance between H3K36me2 and absolute gene expression levels. (b) Gene expression changes in *Nsd2*-edited mice. Significant expression changes in *Nsd2*$^{wt/-}$ and *Nsd2*$^{wt/P906L}$ thymocytes compared to wild type were indicated by arrows based on FPKM. Blue arrows indicate the genes in which H3K36me2 at promoter altered in the same directions as expression changes. Red arrows indicate the genes that did not accompany H3K36me changes at promoter. (c) H3K36me2 decrease and DNA hypo-methylation were more obvious in *Nsd2*$^{P906L/P906L}$ than *Nsd2*$^{wt/-}$ and *Nsd2*$^{wt/P906L}$. (d) Genome-wide DNA methylation changes in *Nsd2*$^{-/-}$ fetal brain and heart. (e) DNA methylation signatures in *Nsd2*$^{-/-}$ fetal brain and heart. (f) DNA methylation changes around transcription start site (TSS) of significantly down-regulated genes in *Nsd2*$^{wt/-}$ thymocytes. (g) DNA methylation changes in gene body in *Nsd2*$^{wt/-}$ thymocytes; significantly down-regulated genes in *Nsd2*$^{wt/-}$ (left) and not expressed genes in thymocytes (right). (h) Proportion of combination of DNA methylation changes and H3K36me2 changes in *Nsd2*$^{wt/-}$ thymocytes by genetic regions. Concomitant decrease of DNA methylation and H3K36me2 at intergenic region was most major changes. (i) *Pcbp3* is one of the genes in which concomitant changes of H3K36me2, gene expression, DNA methylation at TSS were observed in *Nsd2*$^{wt/-}$, *Nsd2*$^{wt/P906L}$, and *Nsd2*$^{P906L/P906L}$ thymocytes.

Fig.S4

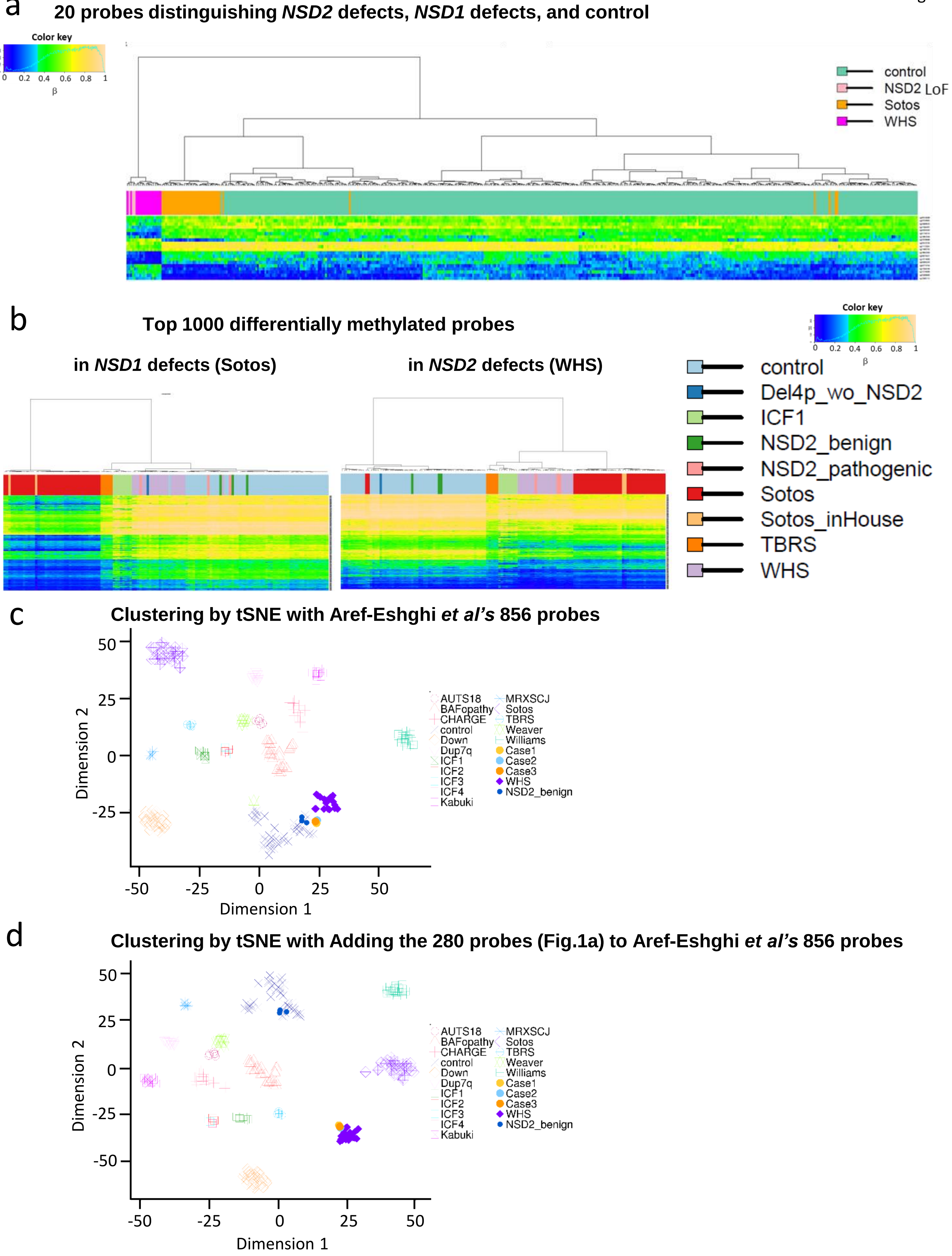
a
20 probes distinguishing NSD2 defects, NSD1 defects, and control
Color key
0 0.2 0.4 0.6 0.8 1
β
control
NSD2 LoF
Sotos
WHS
b
Top 1000 differentially methylated probes
in NSD1 defects (Sotos)
in NSD2 defects (WHS)
Color key
0 0.2 0.4 0.6 0.8 1
β
control
Del4p_wo_NSD2
ICF1
NSD2_benign
NSD2_pathogenic
Sotos
Sotos_inHouse
TBRS
WHS
c
Clustering by tSNE with Aref-Eshghi et al's 856 probes
Dimension 2
Dimension 1
50
25
0
-25
-50
AUTS18
BAFopathy
CHARGE
control
Down
Dup7q
ICF1
ICF2
ICF3
ICF4
Kabuki
MRXSCJ
Sotos
TBRS
Weaver
Williams
Case1
Case2
Case3
WHS
NSD2_benign
d
Clustering by tSNE with Adding the 280 probes (Fig.1a) to Aref-Eshghi et al's 856 probes
Dimension 2
Dimension 1
50
25
0
-25
-50
AUTS18
BAFopathy
CHARGE
control
Down
Dup7q
ICF1
ICF2
ICF3
ICF4
Kabuki
MRXSCJ
Sotos
TBRS
Weaver
Williams
Case1
Case2
Case3
WHS
NSD2_benign

Figure S4. Distinction between *NSD2* loss of function and other syndromes by DNA methylation signatures. (a) Hierarchical clustering of individuals with *NSD2* loss of function (NSD2 LoF or WHS), *NSD1* loss of function (Sotos syndrome 1), and controls by methylation levels of the 20 probes. Heatmap indicates methylation levels. (b) Hierarchical clustering of individuals with *NSD2* loss of function (NSD2_pathogenic or WHS), *NSD1* loss of function (Sotos or Sotos_inHouse), ICF1 with *DNMT3B* variants, TBRS with *DNMT3A* variants, and controls (control, Del4p_wo_NSD2 (negative control), NSD2_benign (negative control)) by methylation levels of the top 1,000 probes whose p-values were top in analysis of linear regression models for methylation changes in *NSD1* defects (left) and *NSD2* defects (right). Heatmap indicates methylation levels. (c) tSNE clustering of individuals with Aref-Eshghi *et al*'s identified probes for discriminations of 14 syndromes (ref 9). These probes were not able to distinguish *NSD2* loss of function from healthy controls by beta values. (d) tSNE clustering of individuals with combination of the 280 probes and Aref-Eshghi *et al*'s identified probes for discriminations of 14 syndromes (ref 9). Addition of the 280 probes made possible to distinguish *NSD2* loss of function from healthy controls by beta values.
